# miR-146a is a Pleiotropic Regulator of Motor Neuron Degeneration

**DOI:** 10.1101/2025.09.04.674013

**Authors:** Dylan A. Galloway, Hunter L. Patterson, Mariah L. Hoye, Tao Shen, Mark Shabsovich, Kathleen M. Schoch, Cindy V. Ly, Timothy M. Miller

**Affiliations:** Department of Neurology, Washington University School of Medicine, St. Louis, Missouri, USA

## Abstract

Amyotrophic lateral sclerosis (ALS) is a progressive neurodegenerative disease affecting motor neurons. Here, we have profiled motor neuron microRNAs (miRNAs) during motor neuron degeneration *in vivo* to gain a better understanding of ALS pathophysiology. We demonstrate that one miRNA, miR-146a, is downregulated in diseased motor neurons despite upregulation in bulk tissue. Genetic deletion of miR-146a significantly extended survival in SOD1^G93A^ mice with heterozygous animals demonstrating the largest benefit. A corresponding reduction in spinal cord gliosis but not motor neuron loss was observed. Finally, we observed that a proportion of miR-146a knockout animals develop spontaneous paralysis, motor neuron loss and chronic neuroinflammation with advanced age. Together these findings demonstrate that a single miRNA influences multiple aspects of motor neuron disease and highlights the complex role for neuroinflammation in ALS pathogenesis.

## Introduction

Amyotrophic lateral sclerosis (ALS) is a progressive, fatal neurodegenerative disease with a lifetime incidence of ∼1 in 400. ALS disease progression is rapid following symptom onset, with muscle denervation, atrophy and accumulating weakness leading to death typically within 3-5 years. Though most cases are singleton without an identified genetic cause, nearly 10% are familial and associated with mutations in key genes (1). Missense mutations in superoxide dismutase 1 (SOD1) were the first causative ALS variants discovered, accounting for approximately 20% of familial and 2% of the total ALS population (2, 3). Transgenic overexpression of mutant human SOD1 in rodents is the most well established ALS model with animals developing progressive motor neuron disease (4, 5), enabling investigations into pathophysiology and novel therapeutics. All disease-modifying therapies currently approved for ALS are efficacious in SOD1 models including riluzole and edaravone (6, 7), as well as tofersen designed to treat SOD1 disease specifically (8).

Neuroinflammation is a pathological hallmark of ALS (9), with non-cell autonomous mechanisms implicated as modifiers or potential instigators of motor neuron disease (10). This is supported by evidence that microglia and astrocytes carrying ALS-linked mutations are capable of potentiating and/or inducing motor neuron degeneration *in vitro* and *in vivo* (11–16). While neuroinflammation is clearly implicated in ALS pathophysiology, the precise molecular underpinnings are still being elucidated with disparate effects depending on the pathways investigated. A current view is that early inflammation confers neuroprotection before gradual replacement by a chronic neurotoxic signature (17).

microRNAs (miRNAs or miRs) are short, noncoding RNAs that regulate gene expression and have been implicated in various biological processes including neuroinflammation. miRNAs associate with Argonaute proteins to form the miRNA-induced silencing complex (miRISC) where the miRNA guides specificity (18). miRISC binding to target mRNAs yields translation inhibition and mRNA decay, ultimately reducing protein output (19). Because the targeting seed sequences of miRNAs are short (6-8 nucleotides), a single miRNA can regulate hundreds of mRNA targets (20). This broad post-transcriptional repression places miRNAs as regulatory hubs during development and disease, making them attractive targets when trying to understand and manipulate gene expression. Previous work from our group and others has demonstrated disease-associated miRNA changes in ALS that are relevant to pathophysiology. For instance, ALS patients and SOD1^G93A^ mice share a set of upregulated miRNAs in the spinal cord and inhibiting a single one of these miRNAs, pro-inflammatory miR-155, extends survival in SOD1^G93A^ mice (21, 22).

Alongside miR-155, miR-146a was initially discovered as an inflammation-induced miRNA and shown to operate in a negative feedback loop to resolve inflammation (23). Transcriptionally activated by NF-kB, miR-146a downregulates several key proteins involved in inflammatory signaling cascades including IRAK1 and TRAF6 (24). Genetic deletion of miR-146a in mice results in enhanced T cell activation, myelopoiesis and autoimmunity (25, 26), but how miR-146a affects motor neuron disease is less clear with both protective and detrimental effects reported (27, 28).

To better understand the role of miRNAs in ALS, we began by leveraging miRNA tagging and affinity purification (miRAP) to track how motor neuron miRNAs change throughout disease in SOD1^G93A^ mice. miR-146a was selected for further investigation due to its consistently lowered expression in motor neurons and upregulation in bulk spinal cord tissue. miR-146a reduction improved disease progression and survival in SOD1^G93A^ mice, corresponding to dampened spinal cord gliosis independent of motor neuron preservation. Surprisingly, we observed that a proportion of aged miR-146a knockout mice developed spontaneous paralysis resembling motor neuron disease, with aged knockout animals having elevated markers of glial cell activation. Taken together, we demonstrate a pleiotropic role for miR-146a in motor neuron disease and highlight miRNA-mediated control of neuroinflammation as a key component of both disease development and progression.

## Results

### Widespread changes in motor neuron miRNA expression accompany disease progression in SOD1^G93A^ mice and highlight miR-146a

We were first interested in understanding how miRNA profiles change in motor neurons over the course of disease in SOD1^G93A^ mice. To accomplish this, we leveraged the miRNA tagging and affinity purification (miRAP) technique (29, 30). Briefly, we crossed mice expressing a Cre-dependent, myc-tagged Argonaute protein (tAgo2) to both ChAT-Cre and SOD1^G93A^ lines, generating ALS model animals in which tAgo2 was specifically expressed in ChAT-positive cells (Chat-Cre; LSL-tAgo2; SOD1^G93A^ mice). By immunoprecipitating Ago2 and its associated miRNAs from homogenized spinal cord, we were able to analyze motor neuron miRNA expression over time in these triple transgenic mice and their littermates lacking mutant SOD1 (Figure 1A). miRNA abundance was analyzed at three different points in disease: a presymptomatic time point (∼70 days), an early disease time point (∼100 days) characterized by hindlimb clasping and the onset of weight loss, and a late disease time point (∼140 days) characterized by paretic gait. Across all time points we observed widespread changes in motor neuron miRNA expression, and these changes collectively became more pronounced with advancing disease progression (Figure 1B, Table S1).

**Figure 1:**
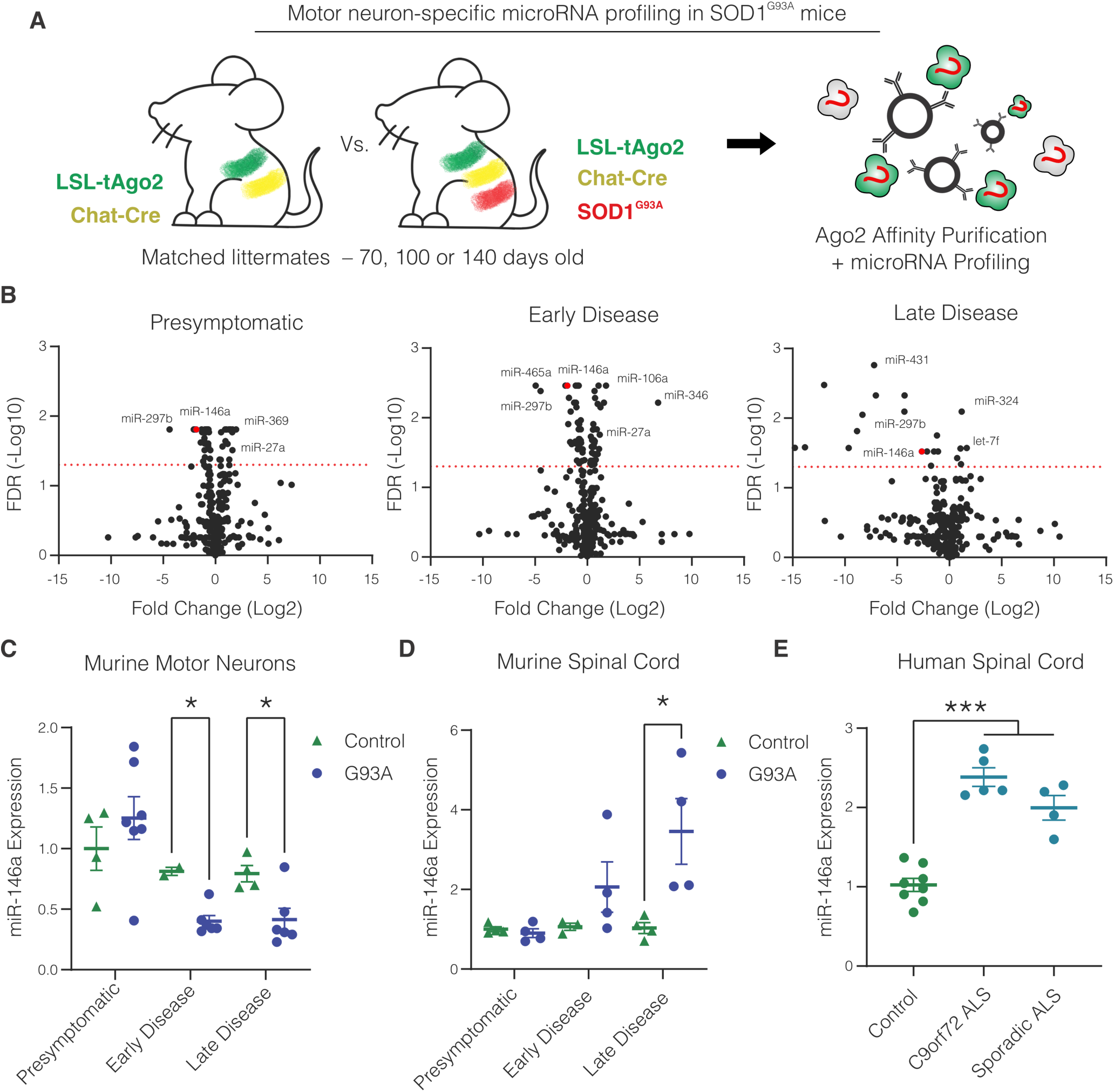
Motor neuron miRNA expression in SOD1^G93A^ mice changes over the course of disease. **(A)** Overview of miRNA profiling in motor neurons from Chat-Cre; LSL-tAgo2; SOD1^G93A^mice collected at presymptomatic, early disease, or late disease time points. **(B)** Fold change was determined by comparing SOD1^G93A^ mice to controls, and the red line marks an FDR threshold of .05. n=3 mice per time point. **(C)** qPCR validation of miR-146a motor neuron expression in an expanded cohort of Chat-Cre; LSL-tAgo2; SOD1^G93A^ mice. Each point represents one mouse. **(D)** miR-146a expression in whole spinal cord from SOD1^G93A^ mice at the same clinical stages (90d,120d, and 160d respectively). **(E)** miR-146a expression in whole spinal cord from human patients without ALS, with C9orf72 ALS, or with sporadic ALS. Mean +/-SEM shown; unpaired t test with Welch’s correction; *p<.05, ***p<.001.

After identifying broad changes, we sought to select a single miRNA to investigate in subsequent experiments. We reasoned that those transcripts exhibiting a consistent trend in altered expression across time points would be more amenable to meaningful manipulation; as in, a miRNA altered in both early and late disease could feasibly modify disease initiation and later progression. We selected miR-146a as a miRNA fitting this pattern and qPCR validation confirmed that motor neuron expression of miR-146a was significantly reduced in SOD1^G93A^ animals in early and late disease (Figure 2C). When measuring miR-146a in whole SOD1^G93A^ spinal cord, we observed a substantial increase with disease progression (Figure 2D). Moreover, measuring miR-146a expression in spinal cord from human patients showed the same pattern. Patients with both C9orf72 ALS, a genetic form of ALS due to a hexanucleotide repeat expansion (31), and sporadic ALS showed double the miR-146a expression of controls (Figure 2E). Analyzing cell type specific profiling in the spinal cord, miR-146a was abundant within microglia and astrocytes (Supplemental Figure 1) (29, 32), that are likely driving the global increase in miR-146a in the bulk spinal cord.

**Figure 2:**
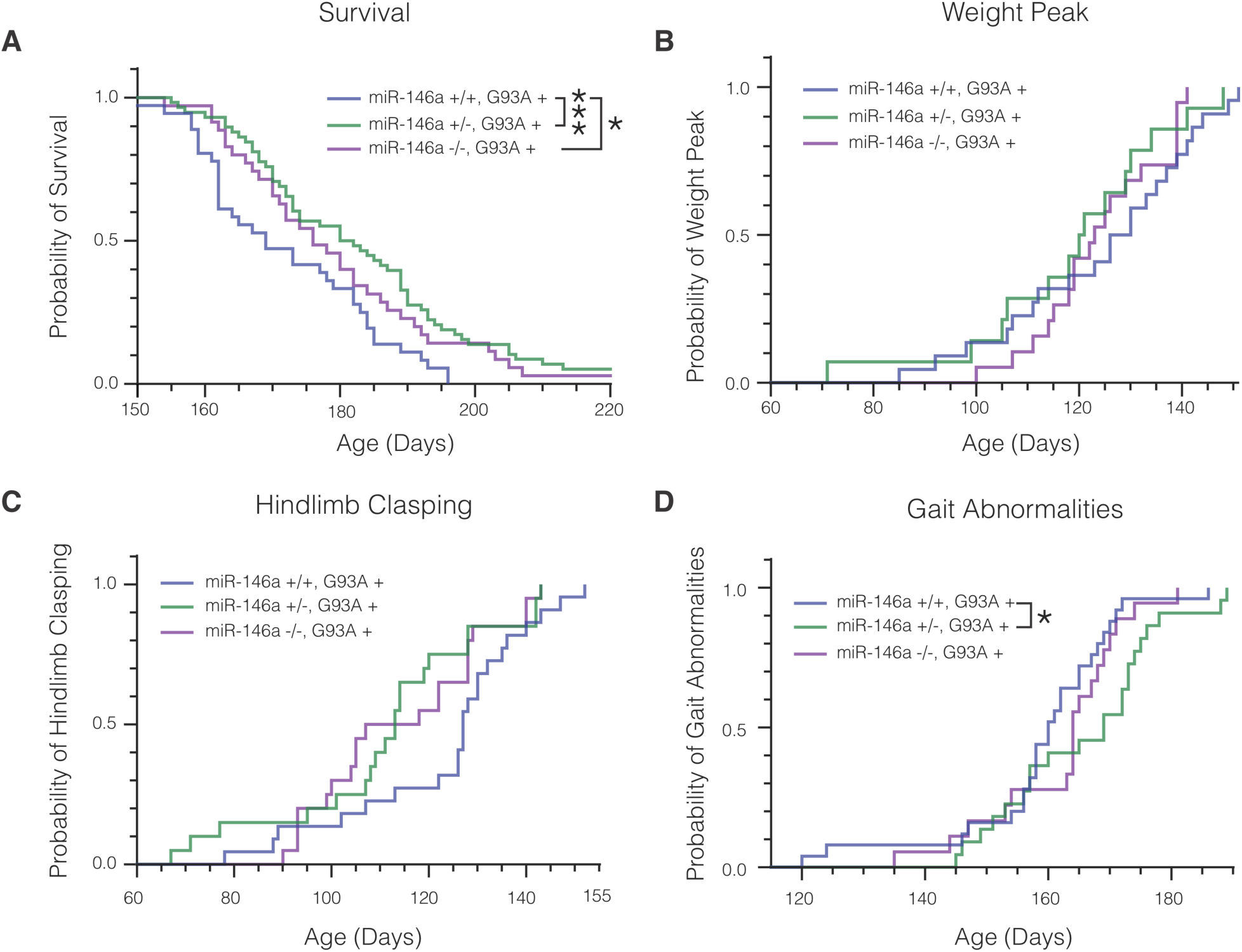
Global miR-146a reduction extends survival in SOD1^G93A^ mice. **(A)** Age at which SOD1^G93A^ mice expressing two, one, or zero copies of miR-146a reached disease endpoint. Median survival was 169 days, 181 days, and 176 days, respectively. N = 36, 58, and 35. **(B)** Age at which mice reached their peak weight. Median age was 128 days, 120.5 days, and 123 days. N = 22, 14, and 19. **(C)** Age at which mice reached NeuroScore 1, characterized by clasping behavior. Median age was 127 days, 113 days, and 112.5 days. N = 22, 20, and 20. **(D)** Age at which mice reached NeuroScore 2, characterized by abnormal gait. Median age was 160 days, 169 days, and 164 days. N = 25, 22, and 18. Log-rank test; * P <.05, *** P <.001.

### Global miR-146a reduction extends survival but does not affect symptom onset in SOD1^G93A^ mice

Given the robust expression changes in miR-146a observed during ALS, we wanted to determine if miR-146a manipulation could affect disease progression. To accomplish this, germline miR-146a knockout mice (26) were crossed to the SOD1^G93A^ line and monitored for survival, weight and behavioral differences (33). We observed that miR-146a reduction, both partial and full, conferred survival benefit for SOD1^G93A^ mutant animals when compared to wild-type littermates (Figure 2A). Heterozygotes survived 11 days longer, on average, where homozygotes showed a milder survival benefit of 7 days. Despite this survival benefit, both heterozygotes and knockouts showed no delay in disease onset as measured by weight loss (Figure 2B) and trended toward accelerated disease onset of about two weeks as measured by hindlimb clasping – though this effect did not reach statistical significance (Figure 2C). Finally, we observed a 9-day delay in the development of paretic gait in mice heterozygous for miR-146a while knockout mice did not show a statistically significant benefit (Figure 2D).

### Global miR-146a reduction limits both astrogliosis and microgliosis in SOD1^G93A^ mice without preserving motor neurons

To determine the mechanism underlying the observed survival benefit in miR-146a deficient animals, we assessed disease pathology at both a presymptomatic (90 days) and late (160 days) disease timepoints. We quantified astrogliosis by measuring the percent coverage of the ventral horn gray matter by GFAP, observing that miR-146a knockouts showed reduced astrogliosis at 90 days while both heterozygotes and knockouts showed reduced astrogliosis at 160 days (Figure 3A, C). Microgliosis was quantified using Iba1, demonstrating reduction in both heterozygotes and knockouts at both time points (Figure 3B, D). To quantify motor neuron loss, we measured the average motor neuron number per ventral horn by counting neurons double positive for both NeuN and ChAT. At both timepoints, there were no significant differences in motor neuron loss across genotypes (Figure 3E, F).

**Figure 3:**
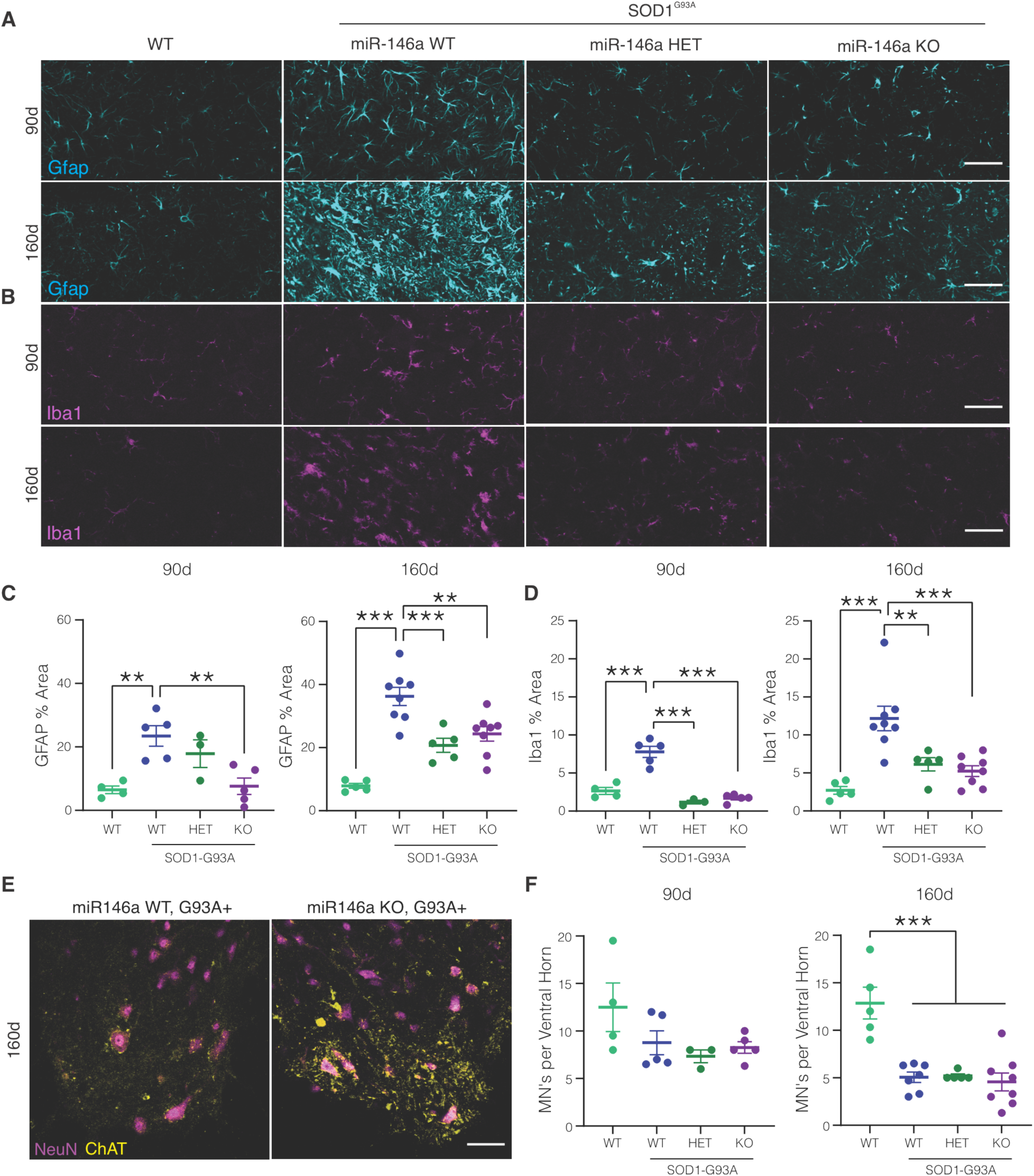
Global miR-146a reduction dampens gliosis but does not preserve motor neurons in SOD1^G93A^ mice. **(A, B)** Astrocytes and Microglia visualized by GFAP and Iba1 immunofluorescence from the ventral horn of mouse lumbar spinal cord. Presymptomatic (90d) and late disease (160d) cohorts are shown for each genotype. **(C, D)** Quantification of GFAP and Iba1 coverage of ventral horn gray matter. **(E)** Representative ChAT/NeuN lumbar cord images used to generate motor neuron counts. **(F)** Motor neuron counts from mice that were presymptomatic (90d) or in late disease (160d). Each point represents one mouse. Mean +/-SEM shown; One-Way ANOVA with Tukey’s test **p <.01, ***p <.001. Scale bars 50 µm

### miR-146a knockout mice show spontaneous paralysis with advanced age

Accompanying our experiments on SOD1^G93A^ mice, we maintained a sizeable number of littermates without the SOD1 transgene to monitor for impairments that could confound SOD1 experiments. No apparent abnormalities presented during the lifespan of a typical SOD1^G93A^ mouse. However, some miR-146a knockout mice began to show sporadic signs of weakness with advanced age (Figure 4A; Supplemental Video 1). This ranged from abnormal gait to hindlimb paralysis and occurred exclusively in miR-146a knockouts. Heterozygous and wild-type littermates did not show this phenotype (Figure 4B). The median onset of paralysis in this cohort was 664 days, at which point half of surviving miR-146a knockout animals had developed some degree of motor impairment. A proportion (∼20%) of knockout animals died without motor symptoms over the experiment consistent with previous reports (26), although we did not observe increased mortality in heterozygous animals in our colony. To better understand the spontaneous disease in these animals, we tracked a subset of animals with biweekly weight and motor assessments. We found that the spontaneous disease was rapidly progressive, with emergence of paralysis, weight loss and death typically within 2 weeks of each other (Figure 4C). In miR-146a KO mice presenting hindlimb deficits, a significant decrease in forelimb strength was also observed (Figure 4D).

**Figure 4:**
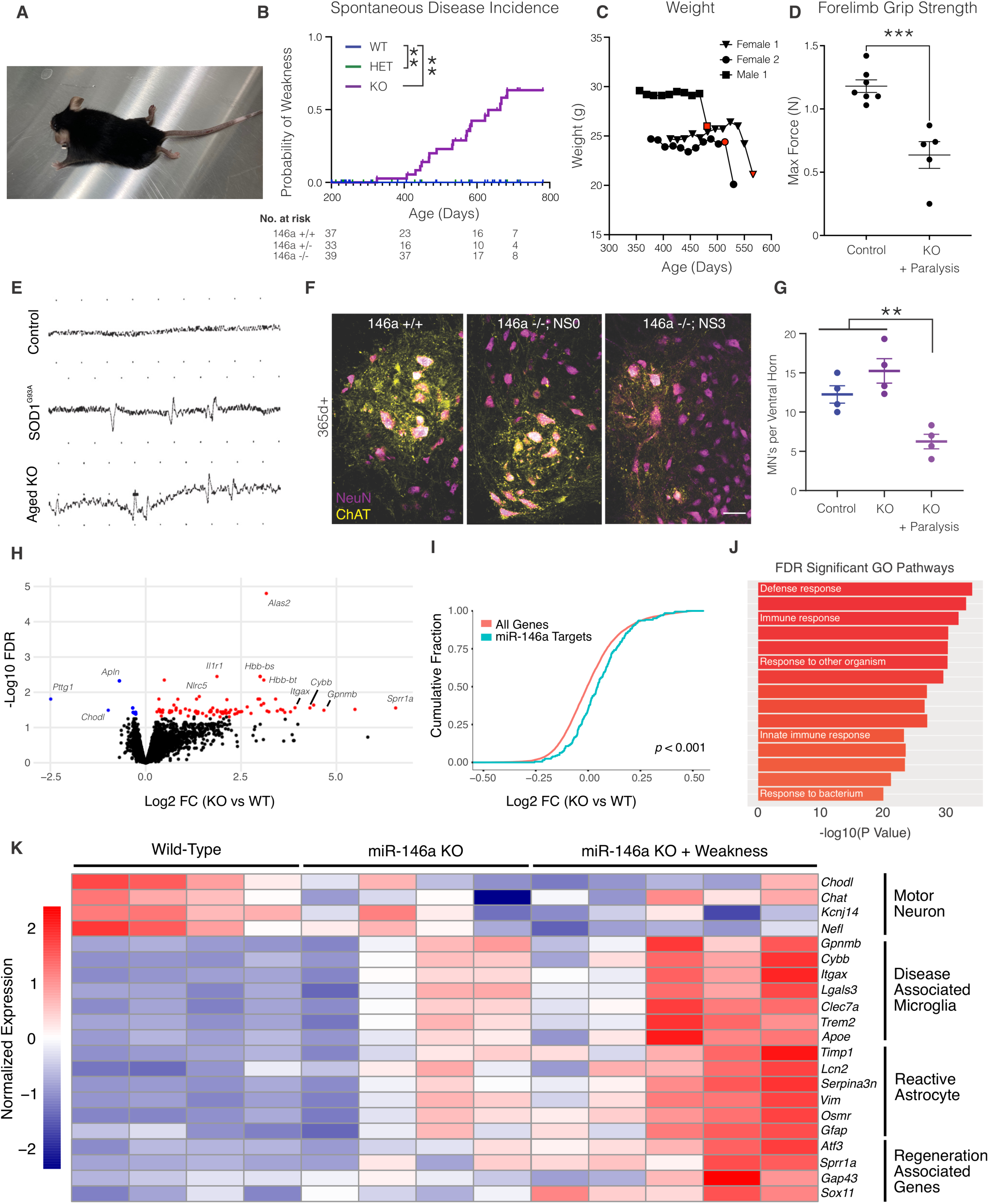
miR-146a knockout mice spontaneously develop motor neuron disease. **(A)** Image of an aged miR-146a KO mouse with hindlimb paralysis. **(B)** Kaplin-Meier curve depicting the cumulative probability of miR-146a KO mice developing weakness. **(C)** Tracked weight from three miR-146a KO mice with red points indicating the onset of paralysis. **(D)** Forelimb grip strength in aged control (miR-146a wild-type and heterozygous animals) and miR-146a KO mice identified with paralysis. **(E)** Needle electromyography in a control, SOD1^G93A^ and miR-146 KO mouse with paralysis. **(F)** Representative ChAT/NeuN lumbar cord images used to generate motor neuron counts. **(G)** Motor neuron counts from control, miR-146a KO or miR-146a KO displaying paralysis. **(H)** Volcano plot of differentially expressed genes in >1 year old wild-type or miR-146a KO spinal cords, colored dots represent genes with FDR corrected *p* values < 0.05. **(I)** Cumulative frequency plot of miR-146a target gene upregulation vs all genes (Kolmogorov– Smirnov test) **(J)** Top significant GO pathways in miR-146a KO spinal cords. **(K)** Gene expression heatmap of select genes. Mean +/- SEM shown; log-rank test **(A)**, unpaired T Test **(D)** and one-way ANOVA with Tukey’s test **(G)**. **p <.01, ***p <.001. Scale bars 50 µm.

Neuromuscular abnormalities were tested in miR-146a KO mice using needle electromyography. We observed an increase in abnormal events, both fibrillations and positive sharp waves, in knockout animals similar to SOD1^G93A^ animals (Figure 4E). Compound muscle action potentials (CMAPs) could not be elicited from the gastrocnemius with sciatic nerve stimulation in miR-146a KO mice but were readily recorded in controls (not shown). Finally, we performed immunohistochemistry for motor neurons in the lumbar spinal cord. We observed a significant decrease in motor neuron counts specifically in aged miR-146a KO animals with paralysis, but not aged control or unaffected KO mice (Figure 4F, G). Thus, the spontaneous disease phenotype observed in miR-146a KO mice resembles motor neuron disease phenotypically, electro-diagnostically and pathologically.

### miR-146a knockout mice show a neurodegenerative glial cell signature in the spinal cord with advanced age

After observing the sporadic development of motor weakness in miR-146a KO mice with advanced age, we explored underlying mechanisms by transcriptional analysis. To assess differential gene expression, lumbar spinal cord transcriptomes from aged control mice and miR-146a KO mice (with and without weakness) were analyzed by RNA sequencing (Figure 4H, Table S2). miR-146a target genes were modestly derepressed in miR-146a KO tissue, confirming a loss of miR-146a function (Figure 4I).

Pathway analysis demonstrated a considerable upregulation of pro-inflammatory gene sets, many of which linked to innate immunity (Figure 4J). Further analysis confirmed that transcripts linked to a common neurodegenerative microglial signature (34) and reactive astrocytes (35) were upregulated (Figure 4K). From a neuronal perspective, we observed *Chodl*, a specific marker of fast motor neurons (36), among the few significantly downregulated genes while several regeneration associated genes canonically associated with axonal injury (37) such as *Sprr1a* and *Atf3* were upregulated. While a set of miR-146a KO animals with paralysis demonstrated the strongest gene expression changes, some miR-146a KO animals without stark weakness showed a similar signature.

## Discussion

In this study, we investigated how miRNA levels change during the course of motor neuron disease *in vivo* and subsequently affect disease pathophysiology. These experiments identified miR-146a as a miRNA that is depleted within motor neurons but upregulated in spinal cord as disease progressed. Genetically deleting miR-146a had pleiotropic effects in SOD1^G93A^ and aging contexts, with miR-146a loss benefiting SOD1^G93A^ animals but predisposing aged animals to motor neuron disease. Given miR-146a’s central role as a regulator of immune responses, our work adds to growing evidence of the Janus nature of neuroinflammation in motor neuron disease.

When investigating miRNA changes in ALS and its models, most previous studies have focused on bulk tissue (21, 22) where motor neurons represent a small fraction of total cells. These studies primarily uncovered upregulation of miRNAs related to neuroinflammation - including miR-155 and miR-146a - suggesting that miRNA measurements in tissue insufficiently capture motor neuron alterations. To overcome this challenge, we utilized an affinity purification method to extract mature, Ago2-loaded miRNAs from motor neurons in the SOD1^G93A^ spinal cord, generating a cell type-specific profile of dynamic miRNA changes during neurodegeneration. Our results indicate that specific motor neuron miRNAs increase or decrease during disease progression, potentially reflecting either cellular adaptations to stress or dropout of specific motor neuron subtypes. These results contrast with the miRNA profiles of human ALS motor neurons generated by laser-capture microdissection (38), where nearly all miRNAs are downregulated indicative of impaired miRNA biogenesis at disease end stage. While we observed many miRNAs downregulated at end stage in SOD1 mice, our results also point to more complex miRNA regulation throughout the entire time course of motor neuron degeneration *in vivo*. As the contributions of many miRNAs uncovered here remain underexplored, we anticipate that continuing to investigate these miRNAs will serve as a window into ALS pathophysiology. While genetic knockout for a large number of miRNAs would be technically challenging, other approaches including antisense oligonucleotide screening (21) or high-throughput identification of miRNA:mRNA interactions (39) could advance our understanding of how miRNAs identified here contribute to motor neuron disease.

Our validation of miR-146a utilized a previously described miR-146a knockout mouse. We generated SOD1^G93A^ animals with heterozygous or homozygous miR-146a deletion and observed extended survival, although disease onset was not delayed. Interestingly, heterozygous animals fared better than homozygotes suggesting that partial reduction of miR-146a is sufficient to impart protection with further reduction becoming detrimental – a finding consistent with complete miR-146a deletion predisposing animals to spontaneous motor neuron disease. Pathologically in SOD1^G93A^ mice, we observed a reduction in gliosis at both early and late time points but not motor neuron rescue. This miR-146a effect mirrors previous work demonstrating that mutant glia drive late disease progression but not onset in SOD1 animals (12), and thus, the observed survival benefit likely results from reduced gliosis in the diseased spinal cord. Why reducing miR-146a, itself a negative regulator of inflammation, limited gliosis in SOD1^G93A^ animals is unclear. A “pre-conditioning” effect is possible, where early enhancement of responses to disease-relevant factors such as misfolded SOD1 or cytokines could stimulate protective immune signaling (40). Another hypothesis is that miR-146a deficiency confers protection via modulation of T cell responses. Several groups have demonstrated that activated T cells have pro-survival benefit in SOD1 transgenic animals (41, 42), and acute activation of microglia by T cell-derived IFNγ is neuroprotective following CNS injury (43, 44). As miR-146a deficient animals have hyperactive IFNγ T cell responses (24, 25), neuroprotective adaptive immunity is an attractive potential explanation for the miR-146a effects observed.

Since miR-146a haploinsufficiency was sufficient to benefit SOD1^G93A^ animals and insufficient to induce spontaneous motor neuron disease, blocking miR-146a activity utilizing antisense oligonucleotides may be a beneficial therapeutic strategy for human ALS. An outstanding question is whether the observed effects of miR-146a stem from the CNS, the periphery or both. As antisense oligonucleotides can be effectively delivered to either compartment (21), future experiments using this approach could both define an effective therapeutic strategy and answer outstanding questions related to miR-146a’s mechanism of action. Other approaches including cell type-specific miR-146a knockout, bone marrow transplants or adoptive transfer of miR-146a deficient cells would be informative, albeit less therapeutically relevant.

During colony maintenance, we observed a small number of exclusively miR-146a deficient mice develop a spontaneous paralytic disease. An expanded aging cohort with multiple measure of motor neuron health confirmed that a proportion of miR-146a knockout animals developed paralysis and motor neuron pathology. The initial description of miR-146a knockout mice reported an elevated incidence of autoimmunity, age-associated cancer and severely reduced survival over ∼1.5 years (26). Our results expand these findings, demonstrating that motor neuron degeneration is another source of dysfunction in these animals becoming apparent with more advanced age. Several overlapping measures of motor neuron health suggest that we have observed a motor neuron disease in our colony, although formally ruling out an autoimmune etiology remains challenging due to low incidence and the requirement for extended aging.

Notwithstanding, our RNA sequencing data confirms a neurodegenerative signature in the lumbar spinal cord of aged miR-146a knockout animals with both microglial and astrocytic markers enriched (34, 35). An interesting note is that a specific marker of fast motor neurons, *Chodl* (36), was one of the few transcripts significantly reduced. Where fast motor neurons preferentially degenerate in ALS mouse models, this vulnerability seems to carryover to miR-146a deficiency. The abundance of genes associated with innate immunity upregulated in miR-146a deficient spinal cords support a model where miR-146a is required to limit excessive neuroinflammation during normal aging, with sustained inflammation sensitizing motor neurons to degeneration. Within the broader NF-κB regulatory network, miR-146a suppresses chronic inflammation by opposing pro-inflammatory miR-155 (45–47), as well as IFNγ signaling (48). As miR-155 is a key driver of the neurodegenerative microglial state (22, 34, 49), removing miR-146a may potentiate the acquisition of this signature via enhanced miR-155 activity. In sum, our studies highlight a critical role for miR-146a as a regulator of motor neuron health and neuroinflammatory responses in the CNS.

## Methods

### Animals

All mice were housed at Washington University in St. Louis under a 12-hour light-dark cycle with ad libitum food and water. Animal procedures were approved by the Washington University in St. Louis Institutional Animal Care and Use Committee (IACUC) in accordance with guidelines from the National Institutes of Health (NIH). Mice expressing tagged Ago2 in a Cre-dependent fashion (RRID: IMSR_JAX:017626), mice expressing Cre in cholinergic neurons (RRID: IMSR_JAX:006410), miR-146a knockout mice (RRID: IMSR_JAX:016239), and mice expressing human SOD1^G93A^ (RRID: IMSR_JAX:004435) were obtained from The Jackson Laboratory to establish our colony. All lines were maintained on a C57BL/6J background. For tissue collection, mice were anesthetized with isoflurane and perfused with 15 mL of chilled phosphate buffered saline (PBS). Tissue samples were flash frozen for molecular analysis or immediately fixed in 4% paraformaldehyde (PFA) for 24 hours if intended for immunofluorescence. After being transferred to 30% sucrose in PBS for at least 2 days, spinal cord samples were frozen in O.C.T. medium (VWR) on dry ice before storage at -80°C.

### microRNA Affinity Purification (miRAP)

Frozen tissue was homogenized and Ago2 pulled down as previously described (30). A BCA assay was performed to normalize input to the lowest protein concentration, and QIAzol (Qiagen) was added directly to the beads for temporary storage and subsequent miRNA extraction.

### miRNA Extraction and Quantification

RNA was extracted from frozen tissue per the manufacturer’s instructions using miRNeasy kits (Qiagen). miRNA microarrays were performed with preamplification using rodent miRNA A+B cards set 3.0 (Life Technologies) on a 7900HT qPCR machine for 40 cycles. To measure the expression of individual miRNA transcripts, reverse transcription was carried out using the miRCURY LNA RT kit (Qiagen) and qPCR was completed using miRCURY LNA SYBR Green reagents (Qiagen) and transcript-specific miRCURY LNA miRNA qPCR assays (Qiagen) on a QuantStudio 12K Flex Real-Time PCR System (Applied Biosystems). miRNA expression was normalized to the geometric mean of previously identified endogenous motor-neuron controls miR-24, miR-30c, and miR-191 (29).

### Animal Behavior

Animals were weighed and behavior was evaluated twice weekly using the NeuroScore (NS) scoring system by two blinded observers (33). Briefly, NS0 was used to define normal mouse behavior, NS1 was characterized by hindlimb clasping, NS2 was defined by paretic gait, and NS3 was defined by bilateral hindlimb paralysis. For survival studies, mice were euthanized when they could not right themselves within 30 seconds of being placed on their sides. The sample size was not predetermined. Forelimb grip strength was measured over five trials using a modified force gauge (Chatillon DFE-II).

### Immunofluorescence and Quantification

Fixed, frozen tissues were cut by microtome to generate free floating 40-micron sections. These sections were washed with tris-buffered saline (TBS) and permeabilized with 0.1% Triton X-100 (Sigma-Aldrich) in TBS. Sections were incubated for 45 minutes in 5% normal donkey serum (Jackson ImmunoResearch) at room temperature to block non-specific binding and subsequently incubated with primary antibody overnight at 4°C. Sections were then washed again in TBS before being incubated with secondary antibodies for 2 hours at room temperature and counterstained with DAPI (Sigma-Aldrich) for 5 minutes at room temperature. Coverslips were applied using Fluoromount-G (SouthernBiotech) mounting media, and images were obtained using an Olympus FluoView1200 scanning confocal microscope. Image analysis was carried out in ImageJ. Motor neuron counts were obtained by counting the number of neuronal cell bodies per ventral horn of the lumbar spinal cord that stained positive for both ChAT and NeuN. For each mouse, this process was repeated on three distinct spinal cord sections and the average number of motor neurons measured was reported. Primary antibodies used include chicken anti-GFAP (Abcam ab4674) at a 1:1000 dilution, rabbit anti-Iba1 (Wako 019-19741) at a 1:1000 dilution, mouse anti-NeuN (Millipore MAB377) at a 1:500 dilution, and goat anti-ChAT (Millipore AB144) at a 1:100 dilution.

### Electromyography

Animals were anesthetized with isoflurane. EMG recordings using a Viking Quest portable EMG machine (Nicolet) were obtained using a 26-gauge, Silicone-coated, concentric needle electrode (Ambu; MFI Medical) and was performed by a certified electromyographer (C.V.L). A 27-gauge subdermal reference needle electrode (Ambu; MFI Medical) was inserted subcutaneously near the recording electrode. A subdermal ground electrode was placed along the midline of the back. The concentric recording electrode was inserted into the tibialis anterior (TA) or gastrocnemius/soleus muscles for recordings. Muscles were evaluated with at least five insertions to assess electromyographic features. Fibrillations and positive sharp waves (PSWs) were identified based on muscle-fiber action potential (MFAP) morphology and regular firing pattern. Nerve conduction studies were performed by stimulating the sciatic nerve near the sciatic notch with constant current square-wave pulses (0.1 ms duration) using a stimulation current increased until a maximal compound muscle action potential (CMAP) could be obtained in control mice then increasing the current by ∼10-20% to provide supramaximal stimulation. miR146a KO mice were stimulated using similar supramaximal current amplitudes used for controls. A recording ring electrode (Natus 39in, 5 Pin) was placed around the gastrocnemius with second reference ring electrode placed ∼1cm distally and subdermal ground electrode placed along the back of the animal.

### RNA Sequencing and Analysis

RNA was extracted from snap-frozen spinal cords using the Norgen Plus RNA extraction kit with genomic DNA removal. Library preparation was performed with 500ng to 1ug of total RNA. Ribosomal RNA was removed by an RNase-H method using RiboErase kits (Kapa Biosystems). Fragmented mRNA was reverse transcribed using SuperScript III RT enzyme (Life Technologies) and random hexamers, followed by a second strand reaction. Fragments were dual indexed and sequenced on an Illumina NovaSeq X Plus using paired end reads extending 150 bases. Reads were aligned to the Ensembl release 101 primary assembly with STAR version 2.7.9a1. Gene counts were derived from the number of uniquely aligned unambiguous reads by Subread:featureCount version 2.0.32. Differential gene expression was performed using edgeR (50), only including genes with > 10 CPM in at least 2 samples.

## Statistical Analyses and Data Availability

Statistical analyses were carried out in GraphPad Prism as detailed in figure legends. miRAP data is found in the data supplement. Sequencing data will be deposited to NCBI GEO upon publication.

## Acknowledgments

We would like to thank members of the Miller lab, Dr. Chris Weihl and Dr. Jony Kipnis for helpful discussions related to the project. RNA sequencing was supported by the Genome Technology Access Center at Washington University in St. Louis. Image acquisition were performed in part through the use of the Washington University Center for Cellular Imaging (WUCCI) supported by the Washington University School of Medicine, The Children’s Discovery Institute of Washington University, St. Louis Children’s Hospital (CDI-CORE-2015-505 and CDI-CORE-2019-813), and the Foundation for Barnes-Jewish Hospital (3770 and 4642).

## Author Contributions

Experiments designed by D.A.G., H.L.P., M.L.H., and T.M.M. Data collection performed by D.A.G., H.L.P., M.L.H., C.V.L. and K.M.S. Animal husbandry and assessment performed by T.S. and M.S. Formal analysis and figure preparation performed by D.A.G. and H.L.P. The manuscript was prepared by D.A.G., H.L.P. and T.M.M. All authors critically revised the manuscript.

## Competing Interests

T.M.M. is a consultant for Ionis Pharmaceuticals, Biogen, and Arbor Biosciences. T.M.M. has licensing agreements with Ionis Pharmaceuticals and C2N Diagnostics.

## Funding

This work was supported by the National Institute of Neurological Disorders and Stroke–National Institutes of Health (Grants R01NS078398 to T.M.M., F31NS092340 to M.L.H.), the University of Missouri Spinal Cord Injury and Disease Research Program (T.M.M.), the Sean M. Healey Center for ALS (D.A.G.) and the Hope Center for Neurological Disorders. The Washington University ALS Postmortem Core is supported by a grant from Target ALS. M.L.H. is currently employed at the National Institutes of Health. This work was completed while M.L.H. was employed at Washington University. The opinions expressed in this article are the author’s own and do not reflect the views of the National Institutes of Health, the Department of Health and Human Services, or the United States Government.

## Supplemental Figures

**Supplemental Figure 1:**
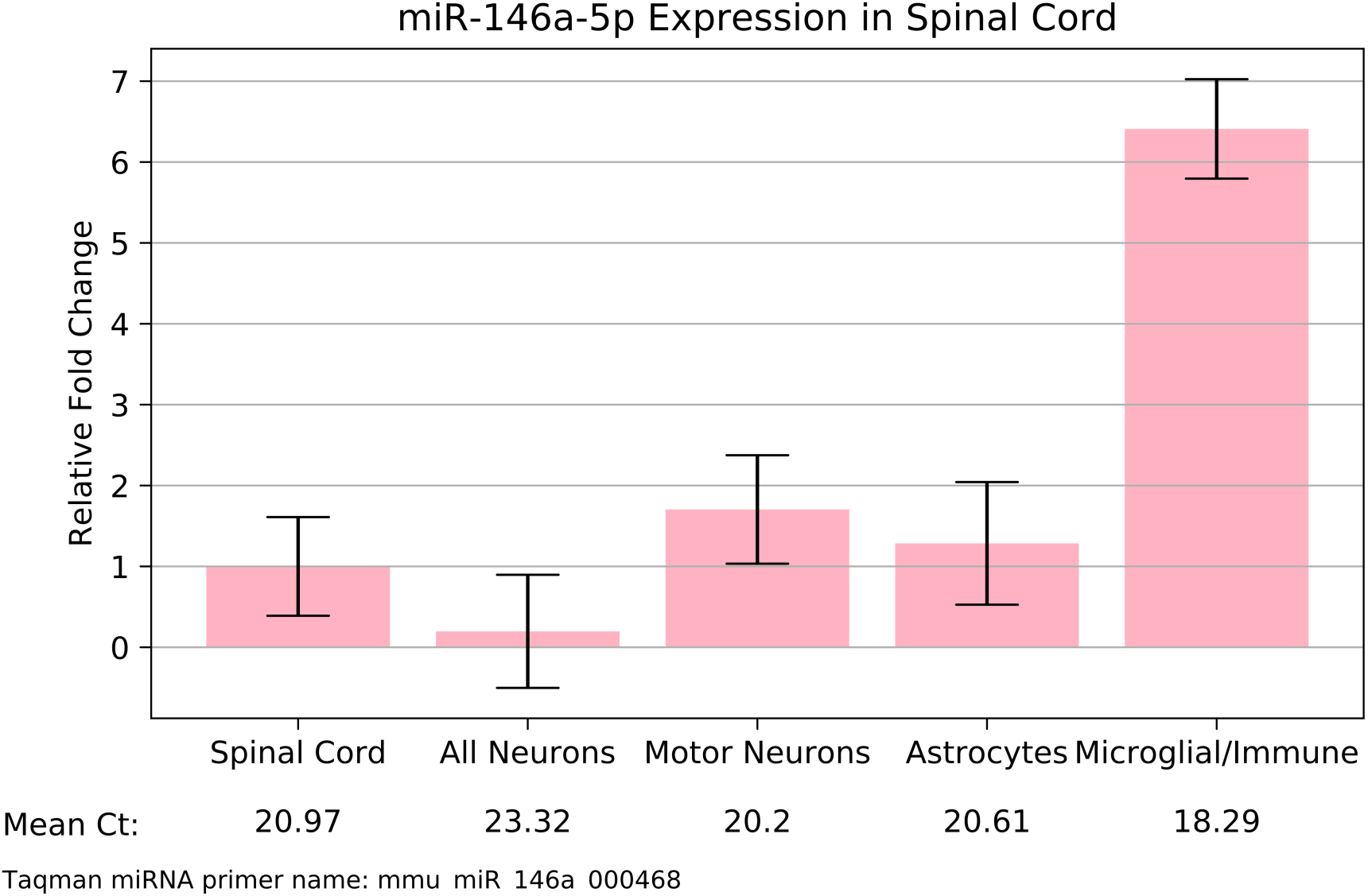
Expression of miR-146a in the mouse spinal cord. miRAP data from miRNA.wustl.edu demonstrating expression of miR-146a in multiple cell types.

